# Diversification, sympatry, and the emergence of mega-diverse tropical assemblages

**DOI:** 10.1101/238329

**Authors:** Jacob B. Socolar, Alexander C. Lees

## Abstract

Geographic gradients in species richness, including latitudinal gradients, can arise from geographic variation in any of three mechanisms: the geologic age of habitats, net rates of evolutionary diversification, or rates of sympatry among diversifying lineages. Here we show that variation in rates of sympatry is a dominant force structuring geographic richness gradients in birds. Species-rich sites contain disproportionately high numbers of recently diverged sympatric species but contain lineages with slower-than-average diversification rates. The positive sympatry-diversity relationship consistently overwhelms the negative diversification-diversity relationship, particularly among high-diversity sites (>250 species). These patterns repeat across biomes and continents with striking regularity, and remain consistent across multiple timescales, including the recent evolutionary past. Biogeographic and evolutionary patterns in birds are consistent with a role for ecological conditions in promoting species coexistence, which allows sister species to co-occur and potentially lowers extinction rates.

## INTRODUCTION

Ecological and biogeographic processes combine to produce pronounced variation in species richness across the Earth, and the mechanisms that underpin geographic richness gradients, including the latitudinal diversity gradient, are a central question in biogeography (Pianka 1966; Ricklefs 1987). At any given location, species richness accumulates due to evolutionary net diversification followed by sympatry of the daughter species. As a result, species richness should be determined by factors that shape regional rates of speciation and extinction, in addition to local processes that govern coexistence. Overall, we might distinguish three broad classes of mechanism that generate richness gradients. *Age-based* mechanisms attribute variation in species richness to variation in the age of a landmass, and thus in the time for diversification to occur (Willis 1922). *Diversification-rate* mechanisms involve geographic variation in rates of net diversification, due either to variation in rates of speciation (Schluter & Pennell 2017) or extinction (Weir & Schluter 2007; Pulido-Santacruz & Weir 2016). *Sympatry-based* mechanisms posit that geographic variation in rates of sympatry drives richness gradients (Pigot *et al.* 2016). Broadly, diversification-rate mechanisms correspond to the evolutionary processes that generate species richness within phylogenetic clades, while sympatry-based mechanisms correspond to the ecological and biogeographic processes that limit the diversity of co-occurring species.

These mechanisms are not mutually exclusive, nor are they independent. For example, sympatry-based mechanisms and diversification-rate mechanisms interact if ecological limits on the diversity of co-occurring taxa (i.e. limits on sympatry) influence net diversification rates (Rabosky 2009; Weir & Price 2011; Kennedy *et al.* 2016; Pontarp & Wiens 2016). Moreover, both mechanisms are required in order to impact species richness. Speciation typically produces daughter species with non-overlapping ranges (Weir & Price 2011; Weber & Strauss 2016), so the subsequent transition to sympatry is therefore a crucial step for the accumulation of alpha diversity at any given location (Pigot & Tobias 2014). Although the different mechanisms interact, we will show that it is straightforward to decompose the contribution of recent diversification to contemporary site-based species richness into separate, multiplicative contributions of net diversification rates and rates of sympatry.

We can additionally distinguish between processes that operated exclusively in deep time versus processes that continue to operate more recently. The boundary between deep time and recent time is arbitrary, and we study timeframes ranging from the past three million years (very recent) to the past fifty million years (relatively deep). For definitional clarity in this paper, we hereafter use *deep time* to refer to processes that occurred more than five million years ago and *recent time* to refer to the last five million years, corresponding to a period when modern bird genera were diversifying (Weir & Price 2011; Jetz *et al.* 2012). The deep/recent distinction is important for two reasons. First, evolutionary patterns and processes may differ between deep and recent time. For example, recent-time diversification rates are reported to be unrelated to bird diversity despite previous analyses suggesting a positive relationship through deep time (Ricklefs 2006; Rabosky *et al.* 2015). Second, mechanisms that operate in recent time can be understood in great detail using present-day ecological and distributional information (e.g. Harvey *et al.* 2017) that is not available for communities through deep time. Thus, if geographic richness gradients arose due to causal mechanisms that continue to operate today, present-day ecology might shed detailed light on the establishment and maintenance of richness gradients.

Here, we investigate the relative roles of diversification and sympatry in structuring geographic richness gradients in birds, using a global avian phylogeny (Jetz *et al.* 2012; Pulido-Santacruz & Weir 2016) and detailed distributional information (Birdlife International & NatureServe 2016). We show that recent-time diversification and subsequent sympatry (hereafter *recent-time processes*) have enhanced richness gradients globally, especially among sites with over 250 species. This pattern universally results from strong sympatry-based mechanisms. Through deep time, we found that diversification-rate mechanisms have attenuated (rather than enhanced) geographic richness gradients. This relationship weakens and becomes more regionally variable in recent time. We found these results to be remarkably consistent across biomes and continents, suggesting that fundamental mechanistic relationships link species richness and rates of sympatry. One intriguing possibility is that a high potential for species coexistence in species-rich areas drives the rapid accumulation of newly-diverged lineages though recent time (Pigot *et al.* 2016), and might also account for negative relationships between species richness and extinction rates (Pulido-Santacruz & Weir 2016).

## METHODS

### The impact of recent-time processes

To assess the contribution of recent-time processes to geographic richness gradients, we considered a counterfactual: what would contemporary gradients look like if recent diversification had not occurred, or if no recently diverged species were sympatric? These two questions are equivalent, because in either case the counterfactual species assemblage at a point is constrained to include at most one species per historical lineage that existed 5 million years ago. Therefore, our question amounts to asking what contemporary gradients would look like if the past five million years of avian diversification were counterfactually “turned off.”

Answering this question is not equivalent to inferring past richness gradients because recent abiotic changes (e.g. glacial cycles) influence contemporary richness patterns (Hewitt 2000) and because extinct lineages might have contributed to past richness gradients in important ways. We seek to isolate the role of lineage diversification and sympatry of recently diverged lineages, not to “turn off” all biogeographic processes that occurred over the past 5 million years. In any case, despite recent innovations (e.g. Winger *et al.* 2014), robust reconstructions of species’ historical ranges are spatially coarse and do not exist for most birds.

We approximate the counterfactual richness landscape by assuming that the avifauna of any point in space would contain exactly the same historical lineages represented in the true contemporary avifauna (Fig. 1). Thus, recent-time mechanisms contribute to the richness of a point in space precisely when sympatry occurs between recently diverged species at that point. We measure the impact of recent-time processes by measuring the degree to which a point’s contemporary avifauna exceeds the richness of the ancestral lineages (as of 5 million years ago) that gave rise to the contemporary avifauna.

**Figure 1.**
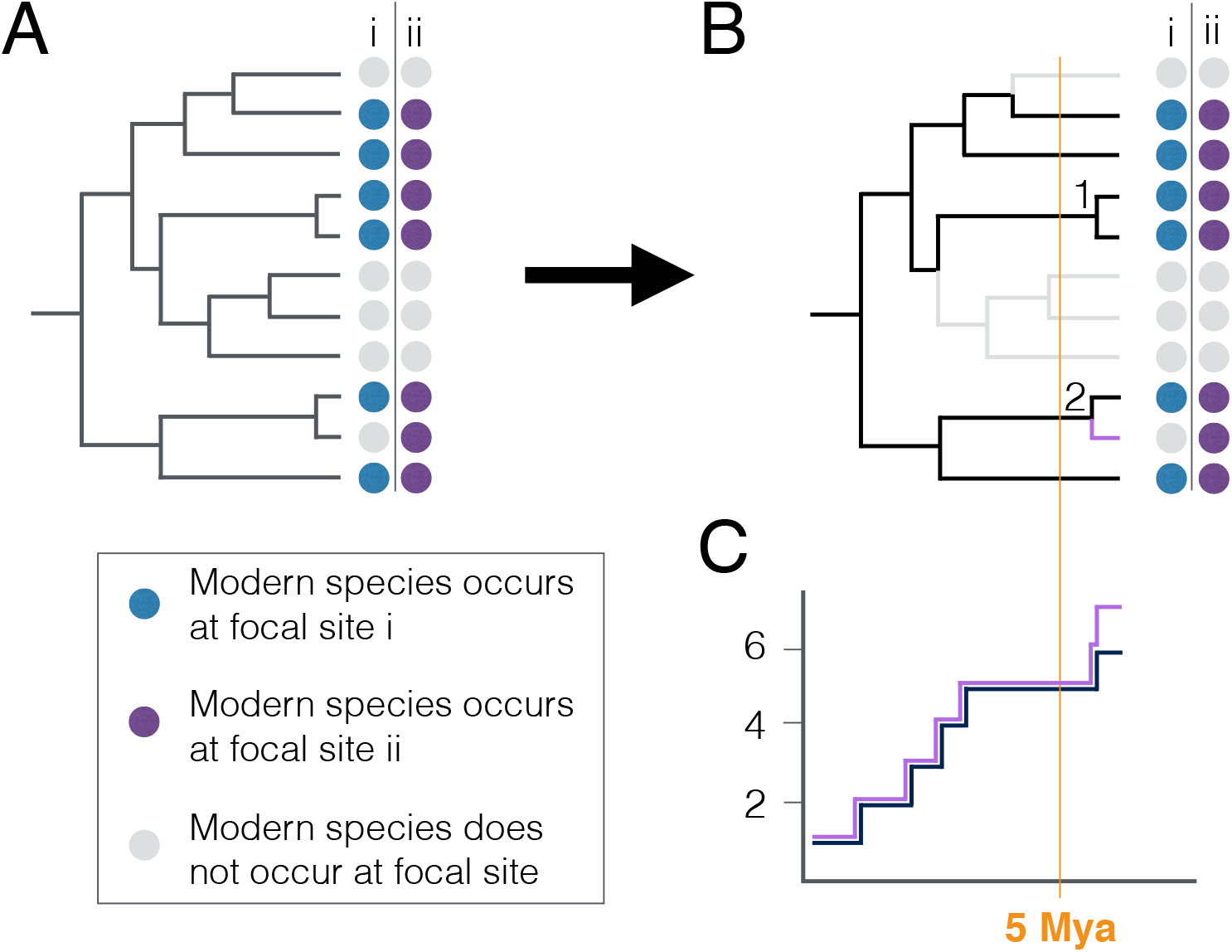
For a given point on Earth (e.g. site i or ii in panels **A&B**), we extract the breeding avifauna **(A)**, prune the phylogeny to contain just those species **(B)**, and examine the accumulation of lineages through time **(C)**. We define the contribution of recent-time processes as the excess of modern species compared to the number of lineages that existed 5 million years ago, expressed as a proportion of the number of historical lineages. Thus, the speciation event at node 1 contributes to the species richness of both site i and ii, but the speciation event at node 2 contributes only to the species richness of site ii.

To quantify the contribution of recent-time processes to species richness, we define **C** to be the set of species that occur at a given location, and **C_R_** to be the number of species in **C** (i.e. the species richness). We then trace the species in **C** back along a time-calibrated phylogeny to find **H** (with richness **H_R_**), the set of historical lineages (as of 5 million years ago) with descendants in **C** (Fig. 1). We define the contribution of recent-time processes to be (**C_R_** - **H_R_**)/**H_R_**. This quantity is interpretable as the degree to which **C_R_** exceeds **H_R_** per historical lineage. Alternatively, it can be re-expressed as **C_R_**/**H_R_** - 1 and interpreted as the degree to which **C_R_** multiplicatively exceeds **H_R_**, corrected to take zero as its minimum value (note that **C_R_** is always at least as great as **H_R_**).

We repeat the process for other time thresholds (other than five million years) ranging from three to fifty million years.

### Distinguishing diversification-rate and sympatry-based mechanisms

There are two ways for **C_R_** to exceed **H_R_** by a large multiplicative factor (i.e. for **C_R_**/**H_R_** - 1 to be large). Either **H** must diversify into a large number of contemporary lineages, or a large proportion of the contemporary lineages derived from **H** must be sympatric at the location of interest. These two situations correspond to a diversification-rate mechanism and a sympatry-based mechanism, respectively (note that age-based mechanisms are unlikely to operate in recent time). We define **T** (with richness **T_R_**) to be the set of all extant species descended from **H**, and we decompose the recent-time processes as follows:

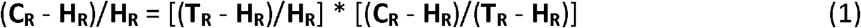

On the right-hand side of equation 1, the first term represents diversification, and the second-term represents sympatry of recently diverged lineages at the location of interest.

### Geographic domain

We restricted our analysis to continental landmasses excluding Antarctica. We excluded oceanic islands because the processes that shape species richness on islands differ systematically from continental processes (MacArthur & Wilson 1967). We included large land-bridge islands of at least 10 million hectares only if they maintained connectivity to the mainland during the last glacial maximum, based on the existence of present-day undersea connections not deeper than 120 meters (Fairbanks 1989; Voris 2000; Jakobsson *et al.* 2008). Most importantly, this resulted in the inclusion of New Guinea within Australasia and the Greater Sundas within Asia (Voris 2000); see Fig. 2). Finally, we excluded large lakes (> 1 million hectares). Smaller lakes comprise a small proportion of the geographic area under study and are not always reflected in the avian range map data that we use. For biome-specific analyses, we used the biome designations in the World Wildlife Fund’s Terrestrial Ecoregions of the World (Olson *et al.* 2001). We lumped humid, dry, and coniferous tropical forests as a single biome, wet and dry savanna as a single biome, and temperate deciduous and coniferous forests as a single biome. Results were generally insensitive to alternative degrees of lumping, though the contribution of sympatry-based mechanisms to the richness of temperate coniferous forests tended to level-off at high species richness (Fig. S1).

**Figure 2.**
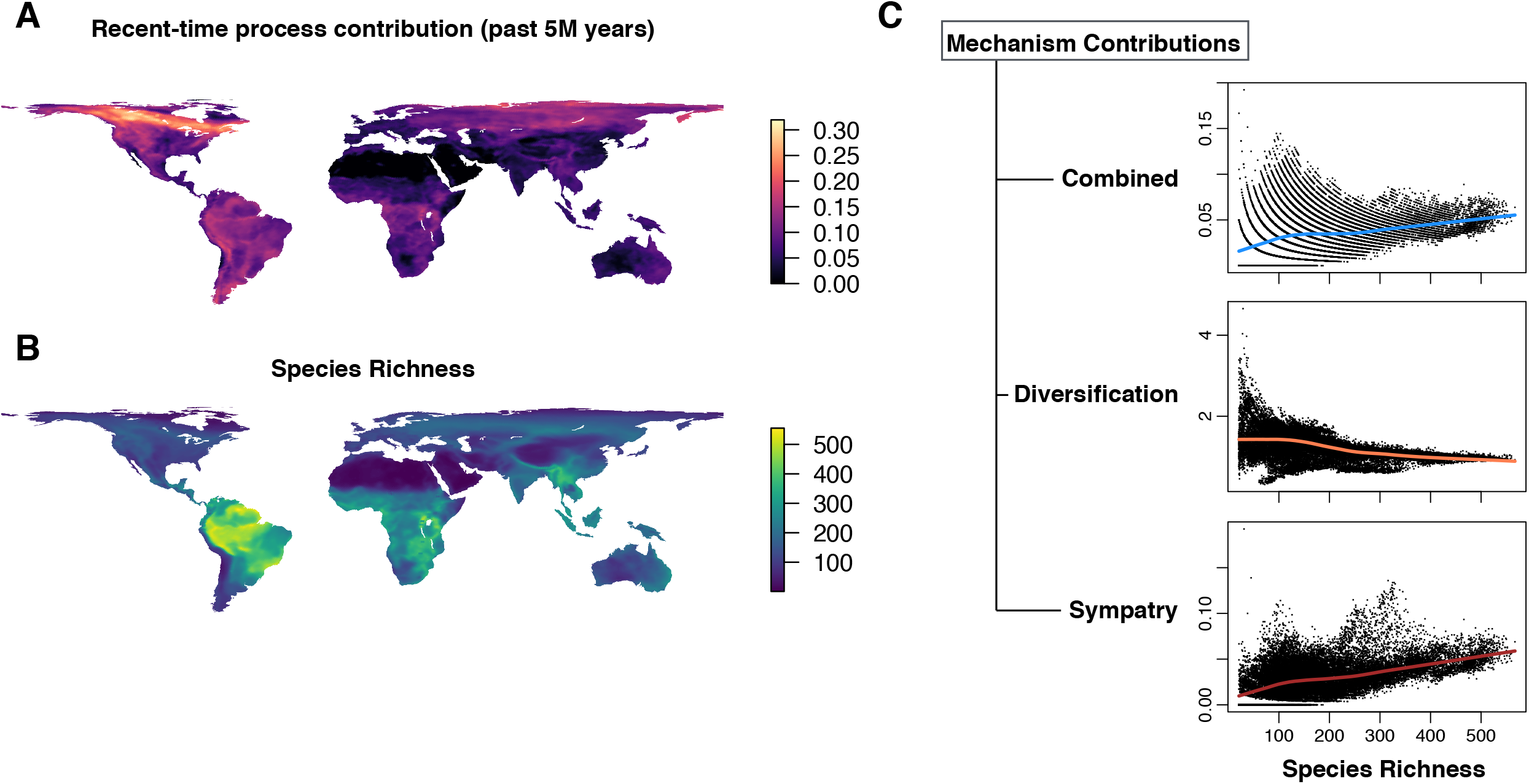
Recent-time processes are positively related to species richness globally, with substantial scatter about the relationship at low-diversity sites. This relationship is the result of a global-scale positive association between species richness and sympatry-based mechanisms that overwhelms a negative relationship between recent-time diversification and species richness. Results are similar across phylogenetic hypotheses (Fig. S4). Numeric values on the y-axes **(C)** follow the quantitative definitions of the methods section.

### Data sources and analysis

We obtained thirty equiprobable phylogenetic hypotheses for extant birds from Pulido-Santacruz & Weir (2016), who updated the global phylogeny of Jetz *et al.* (2012) to include a near-complete molecular phylogeny for the ovenbirds (Furnariidae; (Derryberry *et al.* 2011). We obtained range-maps for each species from BirdLife International (Birdlife International & NatureServe 2016). We standardized species names across the two datasets, combining the range maps for species that are split by BirdLife International but lumped in the phylogeny. Species split by Jetz *et al.* (2012) but lumped by BirdLife were assigned to one of the two relevant taxa in Jetz *et al.* (2012). The BirdLife International maps are the best available global-scale maps for avian breeding distributions. They do not capture micro-scale species sorting due to habitat specialization (Vale *et al.* 2017), and they might overestimate or underestimate the richness of some locations due to decisions about polygon filling, or due to the pervasive assumption of residency in tropical species (Lees & Martin 2015). Nevertheless, these maps have been used with repeated success to study broad-scale trends in avian species richness, turnover, and sympatry (McKnight *et al.* 2007; Jenkins *et al.* 2013; Pigot & Tobias 2014; Cooney *et al.* 2017).

Across a sample of 34,184 equally spaced points over the continental domain of the study (see *Geographic domain*, above), we used the range maps to extract a list of breeding birds. We focused on breeding ranges because the processes governing diversification take place on the breeding range in the vast majority of birds, and because long-distance migration often evolves through shifts in the wintering range, not the breeding range (Winger *et al.* 2014). We then constructed lineages-through-time plots (Nee *et al.* 1994) for the avifauna of each point, and we extracted the richness of both the contemporary species assemblage C and the historical lineages **H** as of 5 million years ago that gave rise to the contemporary avifauna (Fig. 1). For each point, we then calculated the contribution of recent-time mechanisms and we decomposed that contribution into the independent contributions of diversification-rate mechanisms and sympatry-based mechanisms as described above.

We visualized the contribution of recent-time mechanisms to contemporary richness gradients in two ways. First, we mapped the contribution across space. Second, we created scatterplots to display the relationship between overall species richness and recent-time mechanisms, diversification-rate mechanisms, and sympatry-based mechanisms. As a visual aid in pattern recognition, we fit locally weighted smooths (LOESS) to these scatterplots. The fitted curves are useful as visual aids only; spatial pseudoreplication in our dataset precludes statistical hypothesis testing (see *Geospatial considerations*, below). We created these plots at the global level and also separately for major biomes in the Americas, Africa, Eurasia, and Australasia.

Species-poor locations tend to have high variation in diversification contributions due to sampling effects in small communities. For example, a location with only two breeding birds will have a diversification contribution of either 0 (which is minimal) or 0.5 (which is extremely high). To confirm that these sampling effects are not responsible for producing the patterns in our data, we excluded sites with < 20 species from our analysis. Similar results including all sites are available in the supplementary information (Fig. S2).

We repeated the entire process for each of thirty equiprobable phylogenetic hypotheses. All analyses and mapping were performed in the R statistical computing environment (R Core Development Team 2017).

### Geospatial considerations

Many species have large ranges, so spatial pseudoreplicaton complicates inference about the relationship between species richness and recent-time mechanisms. Techniques to mitigate pseudoreplication (e.g. downweighting species with large ranges) necessarily obscure the true geographic relationship between richness and diversification contribution. Therefore, our primary analyses are not statistical. For example, we use Loess smoothing only for pattern visualization, not for statistical hypothesis testing. Instead, we arrive at robust inference based on patterns that repeat with striking regularity across the Americas, Africa, Eurasia, and Australasia. There is every reason to suspect that the past five million years of evolution and biogeography have proceeded independently on these continents. As biogeographers, we may wish for a replicate Earth to test global-scale patterns, but multiple replicate continents are the next best option.

## RESULTS

Across our global sample of 8,108 species at 34,184 equally spaced continental points, the association between recent-time processes and species richness is generally positive, but with relatively high scatter at sites with fewer than 250 species (Fig. 2). Due to the high scatter at species-poor sites, the largest individual point-level contributions from recent-time processes occur at a subset of species-poor sites. Thus, recent-time processes have generally but not universally amplified richness gradients at the global scale, and the pattern becomes more consistent among high-diversity sites.

The overall pattern subsumes stronger but countervailing patterns in diversification and sympatry. Diversification-rate mechanisms are negatively associated with species richness (Fig. 2). Sympatry-based mechanisms, on the other hand are strongly and positively related to species richness (Fig. 2). Thus, in recent time, diversification-rate mechanisms have tended to attenuate global richness gradients while sympatry-based mechanisms have amplified richness gradients.

The global pattern is consistently repeated, and often stronger, across individual biomes and continents. In almost every major biome on every continent, there is a consistent positive relationship between recent-time processes and species richness (Fig. 3, Fig. S3). We illustrate this pattern in detail for tropical forests, where recent-time mechanisms have contributed roughly 15% of the avifaunal richness at mega-diverse sites, and less than 5% at the least diverse sites (Fig. 4). Consistent with the global trend, within-biome patterns are driven by positive associations between sympatry-based mechanisms and species richness that overcome variable (and often negative) associations between diversification-rate mechanisms and species richness. Patterns are similar across the thirty equiprobable phylogenetic trees (Fig. S4).

**Figure 3.**
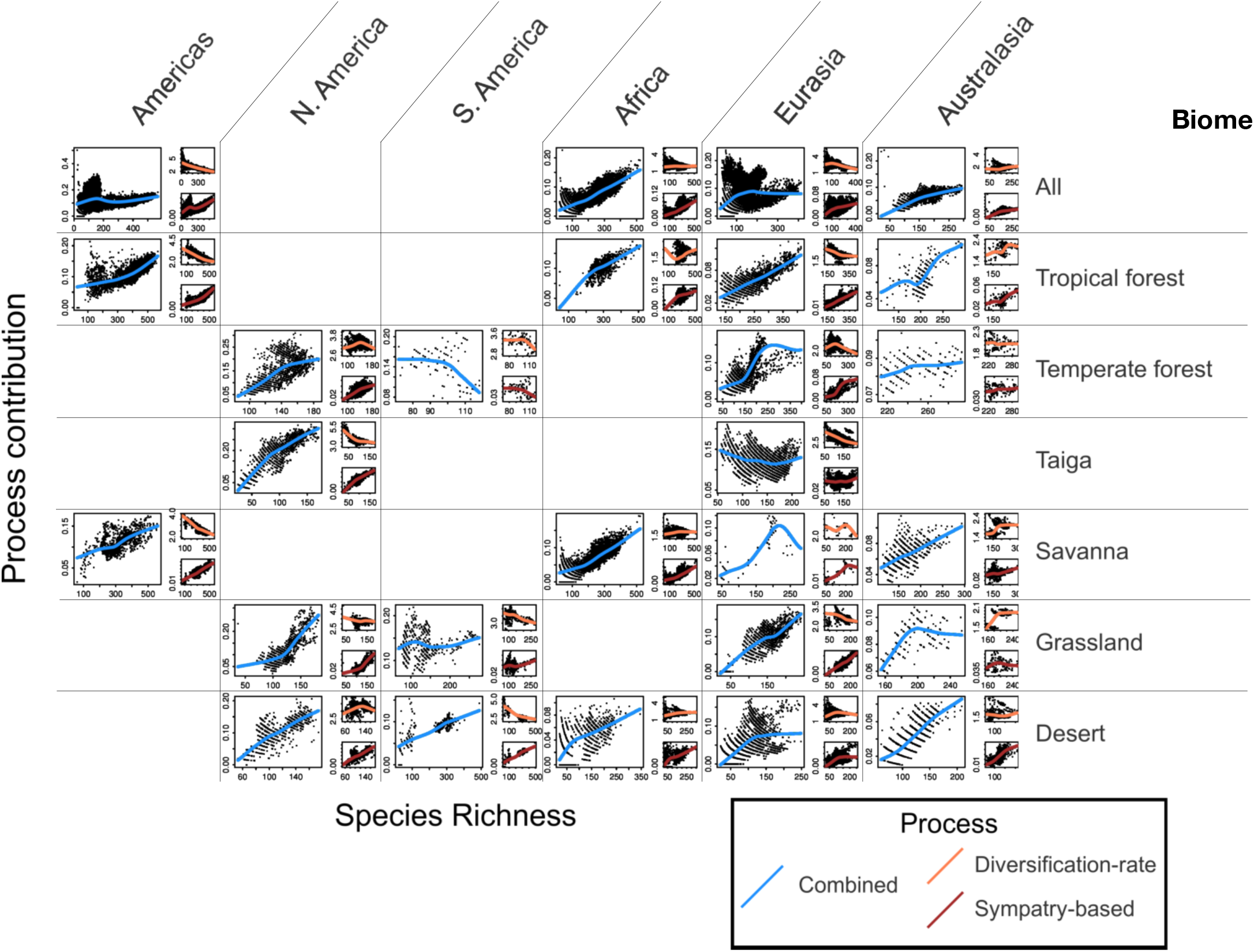
Recent-time processes are positively related to species richness across biomes and processes, reflecting consistent positive associations between sympatry-based mechanisms and richness. Exceptions to this rule mostly involve biomes with small total areas, such as South American temperate forests. Enlarged versions of each panel are available in the supplement (Fig. S3). Results are similar across phylogenetic hypotheses (Fig. S4). Numeric values on the y-axes follow the quantitative definitions of the methods section.

**Figure 4.**
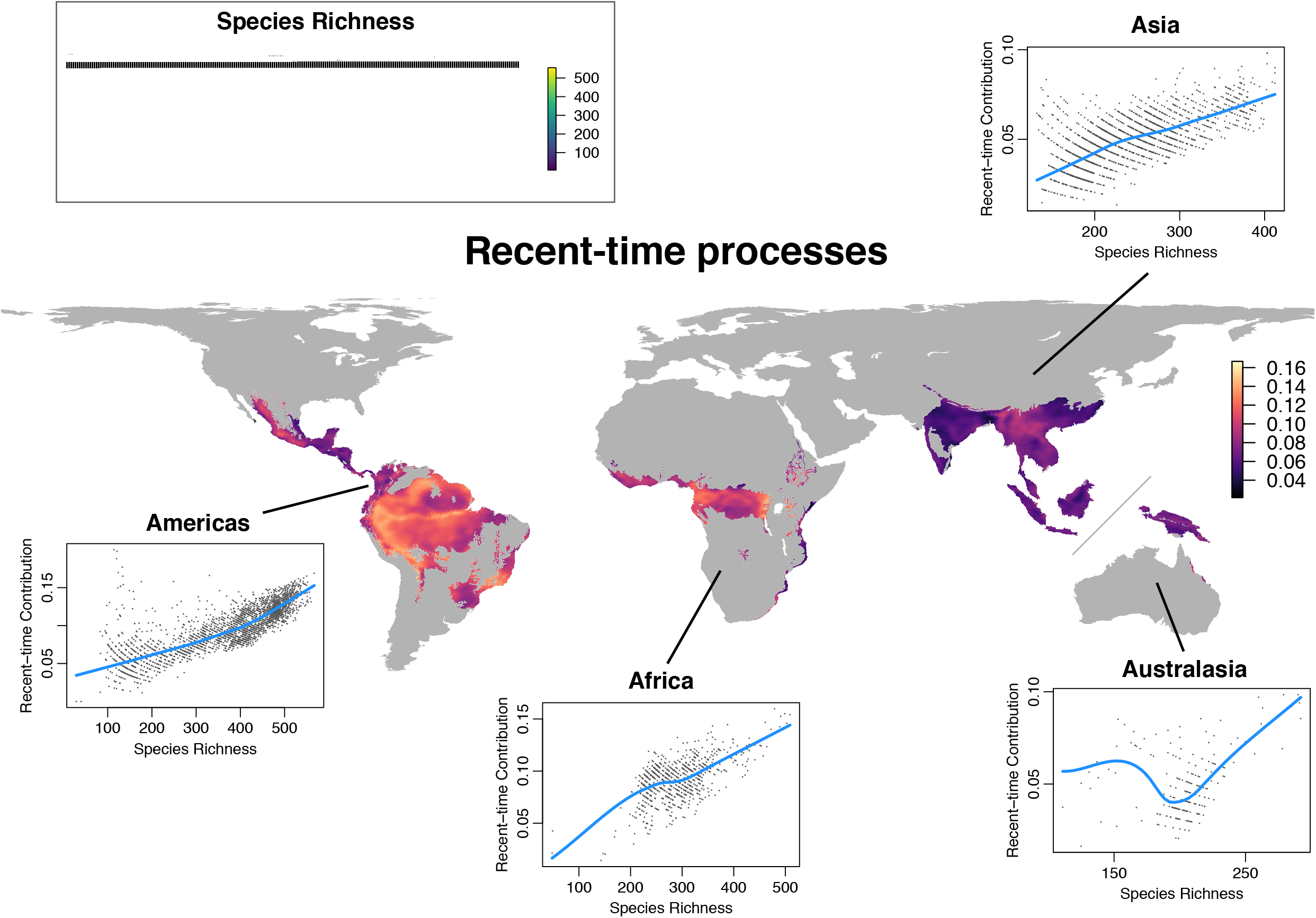
Recent-time processes have contributed strongly to the emergence of mega-diverse tropical forest avifaunas. The richest tropical forest sites derive roughly 15% of their avifauna from recent-time processes, whereas the poorest sites derive less than 5% of their avifauna from recent-time processes. Results are similar across phylogenetic hypotheses (Figure S4).

The patterns described here for the past five million years persist and intensify over longer time intervals, becoming nearly absolute over fifty-million-year intervals (Fig. 5, Fig. S4). Patterns also remain robust through the past three million years (Fig. 5, Fig. S4), though relationships, especially those involving diversification rate, become weaker and more regionally variable.

**Figure 5.**
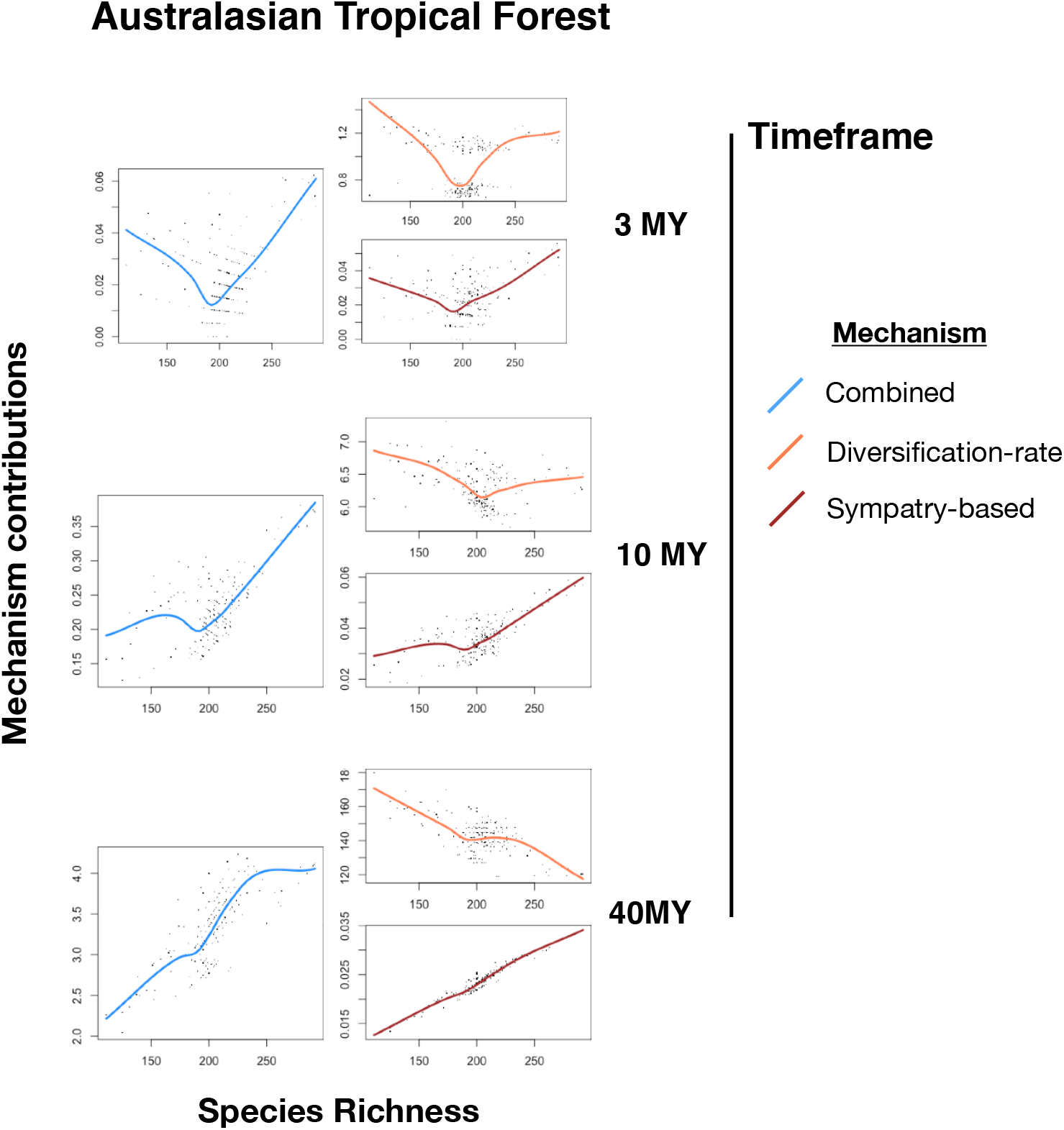
Recent-time relationships remain consistent and intensify over longer time intervals. Shown here are results for Australasian tropical forests. Other biomes show similar patterns (see Fig. S4).

## DISCUSSION

Our framework provides a novel way to understand the contribution of recent avian diversification to gradients in species richness in an explicit geographic context. Previous studies have examined net diversification rates and sympatry separately (Pigot *et al.* 2016; Schluter & Pennell 2017), but both processes must occur together in order for species richness (alpha diversity) to accumulate at any geographic point-location. These processes bear different relationships with species richness, and so neither process in isolation provides a complete picture of how recent evolution has shaped contemporary richness gradients.

Because recent net diversification rates are not associated with latitude, previous work has questioned the importance of recent-time processes in generating species richness gradients (Jetz *et al.* 2012; Rabosky *et al.* 2015; Schluter & Pennell 2017). Our results confirm that no consistent association exists between recent net diversification and species richness, but contrary to previous conclusions we show that recent-time processes have contributed substantially to species richness gradients globally. This pattern is consistent across biomes and continents, and appears to be especially important for the emergence and maintenance of mega-diverse avifaunas. The striking regularity of these results suggests that they reflect robust underlying processes that govern the accumulation of biodiversity in geographic space.

The consistent positive relationship between recent-time processes and species richness arises due to the influence of sympatry-based mechanisms. For example, sympatry among recently-diverged lineages contributes roughly 15% of the species total at the richest tropical-forest sites, but less than 5% at the poorest sites. By quantifying the contribution of recent sympatry to geographic richness gradients, this result builds on the results of Pigot *et al.* (2016), who showed that sister-species co-occur more frequently when they inhabit species-rich regions. The most plausible explanation for elevated rates of sympatry at species-rich sites is the tendency for species-rich sites to permit high levels of species packing, possibly potentiated by high net primary productivity (Pigot *et al.* 2016). Such an effect must be very strong, because it continues to drive recent sympatry at species-rich sites, despite the already-high species richness. Thus, we question the suggestion that low diversification rates at species-rich sites reflect ecological saturation (Schluter 2016).

Diversification-rate mechanisms are often negatively related to species richness in recent time. This result suggests that avian geographic richness gradients cannot be understood by probing the drivers of present-day speciation rates (Schluter & Pennell 2017). Given that the contribution of recent-time processes is generally strongest where net diversification rates are lowest, it appears that recent diversification rates are nowhere low enough to seriously impede the accumulation of new species in local avifaunas. However, recent diversification rates might still play an important role in regulating diversity along habitat gradients that occur at finer spatial scales than the geographic scale examined here. For example, although our analyses show a strong negative relationship between diversification rates and species richness in the Neotropical avifauna, Harvey *et al.* (2017) showed that richness differences across different Amazonian forest types are associated with different rates of recent population differentiation in the species pools specializing on different habitats.

A remaining puzzle is to understand the negative deep-time relationship between net diversification rates and species richness. Over very long time intervals (i.e. stretching back to the common ancestor of all birds), sympatry-diversity relationships as defined here are guaranteed to be one-to-one, and diversification-diversity relationships are guaranteed to be flat. We deal with substantially shorter time intervals in this paper. Based on Ricklefs (2006), our *a priori* expectation was for a positive association between diversification rates and species richness, possibly related to dispersal limitation in speciose avifaunal assemblages (Salisbury *et al.* 2012). We suspect that the disagreement between our results and those of Ricklefs (2006) reflects different choices about which clades to analyze. Ricklefs selected clades of varying ages, some of which are not monophyletic, and regressed clade richness on clade age. We selected all extant clades of a given age (e.g. five million years) and compared clade richness directly. These approaches diverge if, for example, a species-rich clade contains an early-branching lineage that is species-poor. Ricklefs’ (2006) approach requires the researcher to determine whether to treat this as a single species-rich clade, or as two clades of approximately the same age, one species-rich and the other species-poor. Additionally, Ricklefs (2006) compared predominantly tropical clades to predominantly temperate clades, whereas we analyzed all clades represented at a given point and compared points based on their species richness. Finally, Ricklefs (2006) restricted his analysis to passerines, whereas we additionally analyzed the non-passerines.

While we do not advance a mechanistic hypothesis for the negative relationship between diversification rates and species richness, we do note a potential connection between our results and the results of Pulido-Santacruz & Weir (2016), who showed that extinction rates are negatively related to species richness in birds globally. Competition and hybridization can constrain the ranges of recently diverged bird species (Pigot & Tobias 2013), and this constraint might and increase extinction due to elevated extinction risk in small-ranged species or species under pressure from competitively superior congeners that are expanding their ranges. This hypothesis suggests a causal mechanism for the empirically observed connection between species richness, sympatry among sister species, and extinction rates (Pulido-Santacruz & Weir 2016).

The consistency of our results within across space and time strongly suggests that our conclusions our robust to several important caveats and alternative interpretations. First, our measurement of the strength of recent-time mechanisms is relatively unaffected by geographically biased taxonomy (e.g. over-lumping of tropical species), because these taxonomic issues are largely confined to allopatric taxa (Tobias *et al.* 2008). Thus, taxonomic issues are more relevant for studies of species richness in phylogenetic clades than for studies of species richness in geographic space. Moreover, consistent within-biome patterns suggest that geographically biased taxonomy does not fully account for the observed patterns in the decomposition of recent-time mechanisms into diversification-rate and sympatry-based mechanisms (which does depend on patterns of richness in clades in order to calculate **T_R_**).

Second, our assumption that, absent diversification, every point in space would contain exactly the historical lineages represented in the true contemporary avifauna overlooks the role of adaptive radiation in allowing lineages to occupy new environments. If the past 5 million years of diversification have allowed lineages to expand geographically via adaptive radiation, then our assumption might lead us to overestimate the counterfactual richness at some points (e.g. the radiation of *Parulid* wood-warblers in boreal North America). However, our results are consistent across tropical biomes that are thought to be infrequently colonized by radiations from other habitats. The same assumption also overlooks the role of diffuse competition (*sensu* Terborgh & Weske 1975) in constraining species ranges. If diffuse competition from recently diverged, sympatric species constrains the ranges of unrelated species, our assumption underestimates the counterfactual richness of some points. However, an impact of diffuse competition on bird ranges has been observed only in comparisons of highly speciose avifaunas with test localities exhibiting massive species deficits (Terborgh & Weske 1975), and it seems unlikely that marginal increases in diffuse competition due to sympatry between recently diverged lineages would play a strong role in shaping avian ranges.

Third, patterns of richness and sympatry might conceivably arise from edge effects or mid-domain effects, but neither possibility successfully explains patterns globally. Edge effects might include ecotones at biome edges that increase species richness and permit high levels of sympatry via habitat sorting, as well as mountain ranges that act as species pumps (Smith *et al.* 2014) and simultaneously create opportunities for coexistence based on fine-scale elevational segregation. Mid-domain effects might elevate sympatry and species richness if species that evolved in different parts of a biome tend to co-occur near its middle. Both effects undoubtedly play a role generating in global patterns, but neither explains key features of the data. Some biomes (e.g. Neotropical forests) have richness peaks near their middle, while others (e.g. Afrotropical forests) show peaks near their edges. The fundamental consistency of patterns across biomes suggests a single common explanation rather than by local, idiosyncratic explanations. Moreover, Pigot *et al.* (2016) explicitly showed that rates of sympatry among sister species are closely related to gradients in net primary productivity, providing a realistic candidate for the single common explanation.

Global-scale patterns linking diversification, sympatry, and species richness were not previously known to persist through recent time. The recent-time persistence of these patterns raises exciting possibilities for understanding species-richness gradients in terms of the present-day ecology of birds. For example, productivity could potentially allow for high levels of species packing by subdividing the available food resources (MacArthur & Levins 1967), or by supporting longer food chains that allow for species packing on multiple trophic levels and also might promote predation-mediated coexistence within the lower trophic levels (Martin 1988). Alternatively, some feature of species-rich, productive ecosystems could guide evolution along lines that permit rapid sympatry and species packing, for example via sexual selection (Cooney *et al.* 2017) or via foraging traits that are common in productive ecosystems (Salisbury *et al.* 2012). We suggest that teasing apart relationships of this sort might ultimately provide a robust causal explanation for the latitudinal diversity gradient in birds.

## ACKNOWLEDGMENTS

We thank David Wilcove, Alex Washburne, Sam Flake, and the Morgan Tingley lab group for ideas and comments, and Morgan Tingley for support. This work was supported by an NSF GRFP and by the University of Connecticut.

